# QuBiT: A quantitative tool for epithelial tubes reveals cell dynamics and unexpected patterns of organization during Drosophila tracheal morphogenesis

**DOI:** 10.1101/441600

**Authors:** Ran Yang, Eric Li, Madhav Mani, Greg J. Beitel

## Abstract

Biological tubes are essential for animal survival, and their functions are critically dependent on tube shape. Analyzing the contributions of cell shape and organization to the morphogenesis of small tubes has been hampered by the limitations of existing programs in quantifying cell geometry on highly curved tubular surfaces and calculating tube-specific parameters. We therefore developed QuBiT (<u>Q</u>uantitative <u>T</u>ool for <u>B</u>iological <u>T</u>ubes) and used it to analyze morphogenesis during embryonic Drosophila tracheal (airway) development. We find that there are previously unknown anterior-to-posterior (A-P) gradients of cell orientation and aspect ratio, and that there is periodicity in the organization of cells in the main tube. Furthermore, cell intercalation during development dampens an A-P gradient of the number of the number of cells per cross-section of the tube, but these intercalation events do not change the patterns of cell connectivity. These unexpected findings demonstrate the importance of a computational tool for analyzing the morphogenesis of small diameter biological tubes.

## Introduction

Epithelial and endothelial tubes are critical to the survival of almost all multicellular organisms and the physical properties of a tube, including size and shape, are critical to a tube’s function. Misregulation of tube development or maintenance can have severe implications, including polycystic kidney disease in humans (Harris and Torres, 2009; Song et al., 2017). However, the mechanisms by which small tubes control their lengths and diameters is largely unknown (reviewed in (Affolter and Caussinus, 2008; Iruela-Arispe and Beitel, 2013; Wang et al., 2017)). Indeed, even basic descriptions of how cell shape and organization change during development, which is a fundamental prerequisite for understanding size control mechanisms, has not been determined for most tubes.

A major contributor to the lack of characterization of cell shape and organization in tubular epithelia is that existing computational tools for quantifying epithelial cell shape and organization (Barbier de Reuille et al., 2015; Forster and Luschnig, 2012; Khan et al., 2014) are not optimized for tubular epithelia, and in particular for small diameter tubes having high curvature. Moreover, currently available programs do not calculate essential tube parameters such a centerline, branch points, cross-sectional areas and cell orientation. To address this problem, we have created QuBiT (<u>Qu</u>antitation Tool for <u>Bi</u>ological <u>T</u>ubes), a set of computational tools for measuring cell and tube morphology.

As a testbed for developing QuBiT and to identify novel biological processes, we used QuBiT to characterize the early development of the Drosophila trachea system, which is one of the best-studied systems of tubular epithelia (Fig. 1A, reviewed in (Manning, 1993; Samakovlis et al., 1996)). The tracheal system is the gas exchange organ of the fly and thus functions as a lung, but in branch structure it more resembles a vascular system because it is a ramifying network that directly delivers oxygen to specific tissues. Tracheal tubes are epithelial monolayers approximately the size of small capillaries or kidney tubules in mammals, but there are no associated muscle cells, pericytes, or other accessory cells that are known to contribute to tracheal tube size control. Thus, tracheal tube size directly results from interactions of the tracheal cells with each other and with a secreted apical extracellular matrix (aECM) that transiently fills the tube lumens as they expand during their initial development.

**Figure 1.**
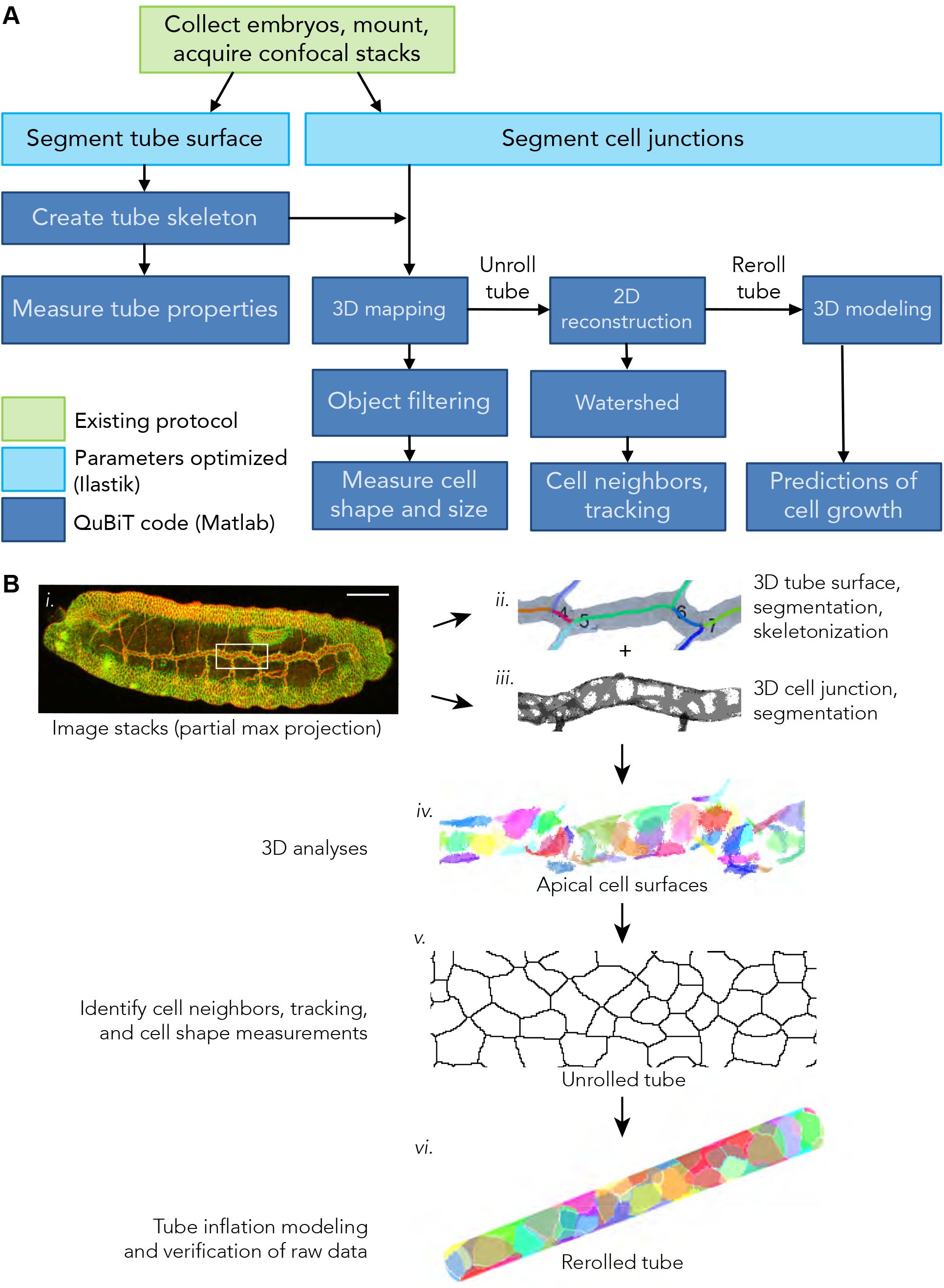
Analysis of high curvature tubes using QuBiT. **(A)** Workflow of QuBiT. Processes that were developed for QuBiT are in dark blue; optimization of existing methods in teal; existing protocol in green. **(B)** Workflow with representative outputs. ***i.)*** A partial maximum projection of a fly embryo at stage 16, with the tracheal system labeled in red with Uninflatable (Uif) and cell boundaries marked in green with septate junction marker Kune-kune (Kune). Scale bar 40μm. 3D outputs for the ***ii.)*** dorsal trunk with calculated centerline and branch points, and ***iii.)*** segmented cell junctions (partial projection). ***iv.)*** Apical surfaces of cells, created using a mask of apico-lateral junctions projected onto the apical tube surface (partial projection). Each cell is individually colored. ***v.)*** 2D unrolling output after watershed segmentation. ***vi.)*** Cells on a model tube after rerolling the 2D output. See main text for details of rerolling.

Using QuBiT, we obtained several unexpected results, including: (1) anterior-to-posterior (A-P) gradients are present in many cellular characteristics, including orientation and aspect ratio, (2) there exists a periodicity at the tube segment level to these characteristics within the A-P gradient, (3) cell intercalation dampens the A-P gradient of the number of the number of cells per cross section of the tube during development, leaving the connectivity distributions of tracheal cells unchanged, (4) cell connectivity distributions in the main tracheal tube are not influenced by the complex shapes of, or possible stresses on, cells that interface the side branches with the dorsal trunk. These results demonstrate both the utility of QuBIT for analyzing small diameter tubular epithelia, and the importance of quantitative analysis in understanding the cell biology of tubular epithelia.

## Results

### Overview of analysis using QuBiT

To maximize maintainability, accessibility, and extensibility of a tool for epithelial tube analysis, we developed QuBiT using commonly available and well supported software platforms rather than develop entirely new programs. The work flow is schematized in Fig. 1A. Image stacks are generated by confocal microscopy using settings that produce cuboidal voxels (Fig. 1Bi). Image segmentation is performed on the entire stack using Ilastik, a general-purpose image segmentation program (Kreshuk et al., 2011). We then analyze segmented images using custom-written code in Matlab (open source available at http://github.com/gjbeitel/QuBiT). Tube analysis proceeds by segmenting the boundary of the tube lumen and creating a skeleton, which enables robust calculations of parameters of interest, including length, surface area, and cross sectional area (Fig. 1Bii, gray tube). Separately, cell junctions are masked onto the tube surface, resulting in apical cell surfaces that can directly be analyzed for parameters such as size and orientation (Fig. 1Biii). While this approach does not yield a full 3-D reconstruction of the entire cell bodies that comprise a tube, it focuses on the apico-lateral junctions and regions that control tracheal cell shape and tube size (Beitel and Krasnow, 2000; Laprise et al., 2010; Sollier et al., 2015; Wodarz et al., 1995), and greatly simplifies the reconstruction problem.

### QuBiT – data collection

QuBiT is designed to analyze images collected with cuboidal voxels containing either tube surface or junctional information with enough resolution for the desired analysis. For example, for the embryonic tracheal system, basic length measurements were performed using image stacks of entire whole-mount embryos collected using a 40X oil objective with a 0.38μm voxel size (Fig. 1Bi). Approximately 75 optical sections were collected per embryo. However, cell parameter analyses required images collected using a 100X oil objective with 0.15μm voxels and the tiling of two image stacks to recreate an entire tracheal tube.

### QuBiT – surface mapping and defining tube centerlines

QuBiT defines a tube surface using markers that either visualize the cell lumenal/apical surface or the lumenal contents themselves (Fig. 1Bi, Bii). For the Drosophila trachea, we used the apical marker Uninflatable (Uif) (Zhang and Ward, 2009) for the analyses presented below, but we have also successfully used the aECM marker Vermiform (Verm) (Wang et al., 2006). We extracted the tracheal system from surface epidermal staining using the Carving module in Ilastik. The segmented data were then imported to Matlab, where we applied a marching cubes algorithm (Cline et al., 1987; Hassouna and Farag, 2007) to define the centerlines of the main and side branches (Fig. 1Bii). The tube centerlines, which are not calculated by other epithelia analysis software, are of critical importance because centerlines define the lengths of tube segments, mark branch intersection points that define tube segments, are the local reference for determining cell orientation, and are necessary for calculating orthogonal planes used to quantify tube parameters including diameter, cross-sectional area and surface area (e.g. Fig. 2C,D). To allow rapid investigation of different parameters without recalculation, computed results from the segmented data are stored with the image stack. For determining tracheal tube parameters, we used 10 cross-sections per segment, which samples the tube approximately once every 15 voxels or 2.3 μm.

**Figure 2.**
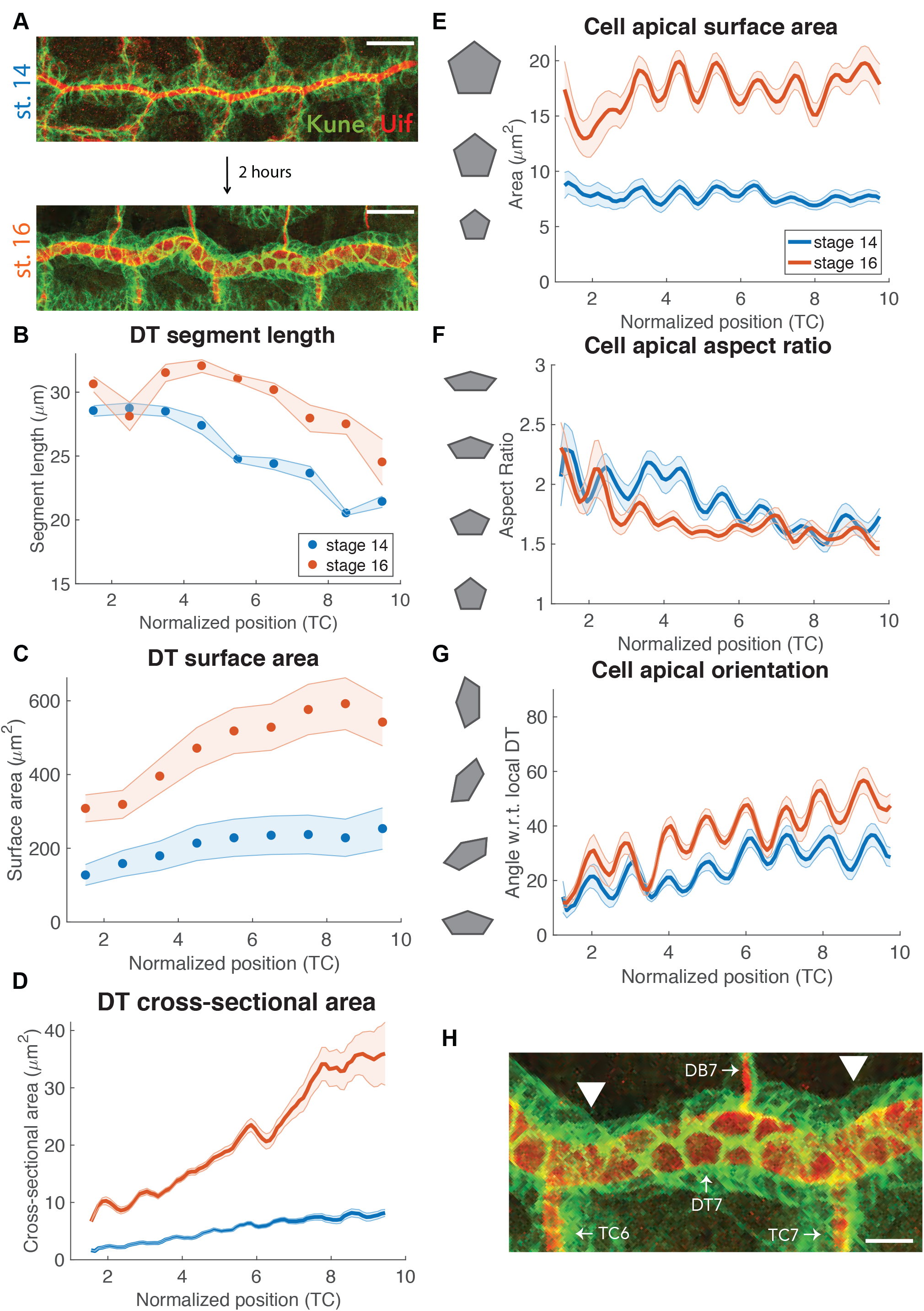
Many parameters of the tracheal dorsal trunk and its cells have anterior to posterior gradients and segmental patterns. Developmental changes between stage 14 (blue label) and stage 16 (orange label) are evident in the shape parameters for the dorsal trunk (B-D) and DT cell apical surfaces (E-G). **(A)** Representative partial extended focus projection images of stage 14 and 16 tubes. Scale bars 15μm. **(B)** Measurements of DT length, by segment. Segment length has a mostly negative anterior-to-posterior (A-P) trend, but the tube grows by 13.7±1.3% in length from stage 14 to 16. **(C-D)** Cross-sections for calculating DT surface area and cross-sectional area were generated by taking projections of the tube in orthogonal planes, resulting in a series of 2D slices that were used to calculate DT surface area in C and cross-sectional area in D. **(C)** Tube surface area, calculated as the average circumference in 10 evenly spaced cross-sections multiplied by segment length, for each DT segment. Within a single tube, surface area increases as a square root function from A-P, and it consistently increases by a factor of 2.2±0.2 from stage 14 to stage 16. **(D)** Tube cross-sectional area. In agreement with surface area, cross-sectional area grows linearly from A-P; and expands by a factor of 4.0±0.1 from stage 14 to stage 16. The steadily increasing diameter expansion ratio is in contrast to the uneven increases in DT length ratio, which is consistent with previous observations that tube length and diameter are regulated independently (Forster et al., 2010; Luschnig et al., 2006; Wang et al., 2006), and further indicate that length is separately regulated in each segment. **(E)** Cell apical surface area increases in a shallow, non-linear gradient such that anterior cells grow by a factor of 1.7±0.2 while posterior cells grow by a factor of 2.6±0.1 (p < 10^−10^, paired t-test between stages 14 and 16). **(F)** We determined cell apical aspect ratio using principal component analysis of the apical surface and found that it has a negative A-P trend at both stages 14 and 16. **(G)** We calculated cell orientation as the angle between the primary component vector and the local vector of the tube centerline. Cell orientation generally increases from A-P and while there is a trend of increased angle at stage 16 compared to stage 14, this trend does not reach statistical significance (p = 0.08, unpaired t-test). **(H)** Higher magnification of the stage 16 image from panel (A). Arrowheads indicate the location of the tracheal segment boundaries and fusion cells. Scale bar, 5μm. For this and subsequent figures using cumulative data, we used six wild-type embryos at stage 14 and six at stage 16, for a total of 692 and 852 identified cells, respectively. Line plots in panels (D-G) were generated using a moving window average with window size of 0.4 normalized tube segments. Dark lines, mean; shaded path under lines, standard error of the mean.

### QuBiT – cell mapping

To map cells onto the tube surface, QuBiT uses information from apico-lateral junctions (Fig. 1Biii). For the tracheal system, we used the claudin Kune-Kune as a marker of septate junctions as we have previously done (Nelson et al., 2010) because Kune staining gives a more continuous signal along the lengths of junctions than the E-cadherin staining, which improves segmentation.

Junctional segmentation was performed using the Pixel Classification tool in Ilastik. We imported this data into Matlab and masked and inverted the junctions on the tube surface to obtain cell surfaces (Fig. 1Biv). We then calculated cell parameters that included area, cell orientation and aspect ratio.

Because the small cell size, narrow diameter and junctional organization of the tracheal dorsal trunk leads to unresolvable segmentation errors, we used several parameters to filter our tracheal cell data. We applied a radial filter to exclude cells on dorsal branches and transverse connectives, an axial position filter to remove cells outside of regions of interest on the dorsal trunk, and a size filter to exclude improbably small or large cells (Fig. S1A).

### QuBiT – Re-imaging: unrolling the tube to make a 2D surface model

To provide a planar representation of the cells that allows direct application of a broad array of existing 2D analysis tools, QuBiT can computationally unroll and flatten tubes. To do this, we calculate the exact cross-section using the orthogonal plane to the local centerline at regular voxel-sized intervals along the length of the dorsal trunk. We then extract the cell junction data from projections radiating from the centerpoint in each orthogonal plane and write the data on a 2D plane (Fig. 1Bv).

As an example of an operation that is much easier to perform in 2D, we used watershed segmentation to reduce cell junction widths to single pixels (Fig. 1Bv). This enables analysis of parameters such as cell connectivity (the number of cells any given cell touches) and the number of cells in a tube cross section, which we utilized in the analysis of the tracheal system below. In future work, 2D projections could be used to track tracheal cells through time more easily than in 3D, which is important in determining changes in cell arrangements.

### QuBiT - Re-imaging: recalculating and rerolling a 2D projection into a 3D model

The process used in QuBiT to unroll a tube can also be utilized to recreate a tube from a 2D projection (Fig. 1Biv). As an example of how this can be employed, we used tube unrolling and rerolling to test for evidence of active cell rearrangements during tracheal expansion. We computationally expanded embryonic stage 14 tracheal tubes to the size of stage 16 tubes, and then compared the 3D parameters of the computed and actual stage 16 apical cell surfaces.

Tubes were computationally expanded by projecting the raw-unrolled 2D stage 14 cell data back onto 3D tube surfaces with corresponding lengths and radii of stage 16 tubes. To do this, we increased the number of pixels along the long (X) axis of each 2D projection proportionally to our measured changes in tube length using nearest neighbor interpolation. Then at each position along the model tube axis, we sampled the 2D image for cell surfaces and wrote the data on the orthogonal (YZ) circumference with corresponding radius based on stage 16 tube radii, which was normalized with respect to axial position. This resulted in a re-creation of stage 14 cell surfaces after tube inflation to stage 16 with no other changes to cell geometry or topology. One limitation of the resulting model is that the projected tube is perfectly straight, whereas the original tubes had small irregularities, which introduces a small amount of error. However, because of the asymmetric expansion along the length of the tube, the alternative approach of expanding along a curved path would result in over and or under expansion of some regions, which would also result in slight deviations. To determine whether systematic errors were introduced, we compared the effect of unrolling and rerolling stage 14 and 16 tubes (Fig. S2). For both stages, there was excellent agreement between the original and rerolled tubes, with most parameters differing by no more than 6%, which is sufficient for the analyses in described below.

### QuBiT – data analysis

For statistical analysis of tube parameters, QuBiT takes advantage of the extensive tools of the MATLAB′s framework to allow users to perform statistical analyses on individual tubes, and also on multiple independent tubes simultaneously, without the need to export the data into other analysis packages. Notably, for tubular systems with reproducible features, QuBiT has the ability to track, align, and compare identifiable features. In the case of the trachea, we aligned individual dorsal trunk segments rather than just normalizing to length along the tube. As detailed below, this enables calculations that reveal tracheal cell parameters differ not just between segments, but also within segments.

### Tracheal dorsal trunk dimensions change in anterior-posterior gradients, but posterior is not always bigger

We tested the utility of QuBiT by investigating in wild-type embryos, the growth of the tracheal dorsal trunk (DT) that occurs during ″tube inflation″, a ~2.5 hr event during which a burst of lumenal secretion (Tsarouhas et al., 2007) doubles tube diameter and increases tube length by about 15% between stages 14 and 16 (Fig. 2A). Our measurements through QuBiT show that the dorsal trunk length increased by 13.7±1.3% (Fig. 2B). These data were not statistically different from our manual length measurements of the same embryos (250±18μm by hand in Volocity vs. 263±6.9μm using QuBiT at stage 16, p = 0.17, two-sample t-test), but had the advantage of automatically generating measurements on each individual segment that allowed a more detailed analysis of the sizes growth of tracheal tubes than had previously been performed. For example, while one might have expected the individual dorsal trunk segments to all expand equally, most of the dorsal trunk growth resulted from increased length of segments DT4 through DT9 (+22±8%), while the remaining segments lengthened only slightly (Fig. 2B). Interestingly, anterior segments are generally longer than posterior segments, despite anterior segments having fewer cells (e.g. ~21 in DT3 vs. ~28 in DT9 (Robbins et al., 2014)). These results predict that anterior and posterior cells have either different arrangements and/or shapes in their respective segments, which, as described below, is indeed the case.

### Many dorsal trunk cell properties have an anterior-to-posterior gradient

To investigate how the shapes of the tracheal cells might contribute to control of tube size, we measured shape parameters of tracheal cell apical surfaces at stages 14 and 16 (Fig. 2A). We focused on the apical surface because the available evidence indicates that tracheal size is regulated at the apical cell surfaces (Laprise et al., 2010; Nelson et al., 2012; Olivares-Castineira and Llimargas, 2017; Robbins et al., 2014), and that the shape of basolateral surface is not regulated (Beitel and Krasnow, 2000).

We first examined how the apical area of individual tracheal cells changes as a function of segmental location and stage. Before tracheal tube expansion at stage 14, tracheal cells in all segments have similar apical areas (Fig. 2E, blue line), and though there is some variability within each segment, there is no apparent A-P gradient. During tube expansion, the area of all tracheal cells increases (p < 10^−47^, comparing stage 14 to stage 16, paired t-test), but posterior cell area increases somewhat more, resulting in a shallow A-P cell apical area gradient (Fig. 2E). This shallow gradient is in marked contrast to the much more pronounced and uniform A-P gradient in anterior vs posterior segment surface area where the A-P difference is almost 2-fold (Fig. 2C). The discrepancy between the fairly uniform cell area and graded segment surface area predominantly results from posterior segments being both shorter (Fig. 2B) and having more cells than anterior cells (~21 cells in DT3 vs. ~28 cells in DT9) (Samakovlis et al., 1996).

We next investigated apical cell shapes by calculating their aspect ratios (Fig. 2F). In contrast to the mostly uniform cell area of tracheal cells, the shape of tracheal cells is not uniform. Anterior cells have higher aspect ratios than posterior cells (p < 10^−3^ at both stages 14 and 16, unpaired t-test of DT2 vs DT10), but overall, aspect ratio does not change significantly during development (p = 0.21, paired t-test).

As with cell size and aspect ratio, cell orientation also shows an unexpected A-P gradient (Fig. 2G). It was previously shown from an analysis of a small number of cells that the long axes of tracheal cell apical surfaces in tracheal segment 8 (DT8) are not aligned with the long axis of the tracheal tube, but instead lie at an approximately 42±21° angle at stage 16 on average (Forster and Luschnig, 2012; Nelson et al., 2012). In excellent agreement with these previous measurements, we found that the average cell orientation in DT8 was 37±12°. However, when we measured cell orientations in all tracheal segments, we found that anterior cells tend to be oriented along the trunk axis with an average angle of 28±5°, whereas posterior cells tend to be oriented along the circumference of the tube with an average angle of 46±4° (unpaired t-test using DT2 against DT9 at stage 16, p < 10^−4^). The gradient exists at stage 14 (p < 10^−3^) and cell orientations differ significantly between stages 14 and 16 (p < 10^−14^, paired t-test).

### Many dorsal trunk cell properties have segmentally repeating variations

Despite tube surface area increasing fairly linearly from anterior to posterior (Fig. 2D), apical cell areas and aspect ratios at stage 16 show a strongly periodic pattern, with cell area and aspect ratio having local minima close to the points where the transverse connectives join the dorsal trunk, and local maxima at the middle of a segment (Fig. 2E, 2F). Cell orientation also shows a sinusoidal pattern, but the phase is opposite to those of surface area and aspect ratio, with orientation having local maxima where the transverse connective (TC) branches connect to the DT (Fig. 2G). This sinusoidal pattern is robust with respect to the window size used to calculate average cell apical surface area along the tube, with the periodicity being clearly visible with window sizes ranging from at least 0.6 to 0.3 of a segment (Fig. S1A). Notably, the sinusoidal pattern corresponds to the physical organization of the trachea. The dorsal trunk is comprised of nine distinct segments that derive from nine clusters of epidermal cells during development. Rather than the tube being a continuous cobblestone of cells, the cobblestone pattern is punctuated by pairs of thin, washer-like cells called ″fusion cells″ that join adjacent segments just posterior to where the transverse connectives branch from the dorsal trunk (Fig. 2H, arrowheads). DT cells close to the fusion cells have reduced apical sizes relative to DT cells more distant from fusion cells.

One explanation for this segmental sinusoidal periodicity could be that the fusion cells constrain tube growth such that the tube is wider between fusion points. However, WT stage 16 DT tubes do not show constrictions at the fusion cells (Fig. 2A, H), and the plotted measurements of tube cross-section area and radius do not show segmental periodicity (Fig. 2D). Thus, the differential areas of cells close to or further away from fusion appear arise from differences between the cells and/or difference in the local lumenal environment. As no periodicity has been observed in lumenal proteins or organization, it then seems likely that the DT cells closer to fusion cells expand their surface area less in response to secreted lumenal contents than do DT cells in the middle of the segment. Consistent with this hypothesis, uniform overexpression of the glycoprotein Tenectin in the DT using the *btl*-Gal4 driver results in tracheal segments in which the diameter of the tube increases more between fusion cells than at fusion cells, with the change in diameter along the segment being a smooth curve rather than a step at the fusion cell (Fig. 6F in (Syed et al., 2012)).

### Tracheal cells maintain cell shape parameters by reorganizing during tube expansion

Given that expansion of the tracheal length and diameter is highly asymmetric, with length and diameter increasing by 13% and 100% respectively, we asked if the changes in cell size, aspect ratio, and orientation could result simply from an inflation of the tube, as if one were to inflate a cylindrical balloon with cell apical surfaces drawn on the surface. This is an appropriate model for the tracheal system because there is no change in cell number during tracheal development (Samakovlis et al., 1996); therefore changes in tube size must result from changes in cell shape and/or cell arrangement. As described above, we used QuBiT to computationally expand stage 14 tubes, and consequently the cells that make up the tube, to their respective stage 16 sizes (Fig. 3Aiii). We then analyzed the cell shape and orientation parameters of the resulting ″computationally expanded″ cells (3Aiii) and compared them to cells from stage 14 and to 16 tubes that had been similarly unrolled and rerolled, but without computational expansion (3Ai and 3Aii, respectively, termed ″model″).

**Figure 3.**
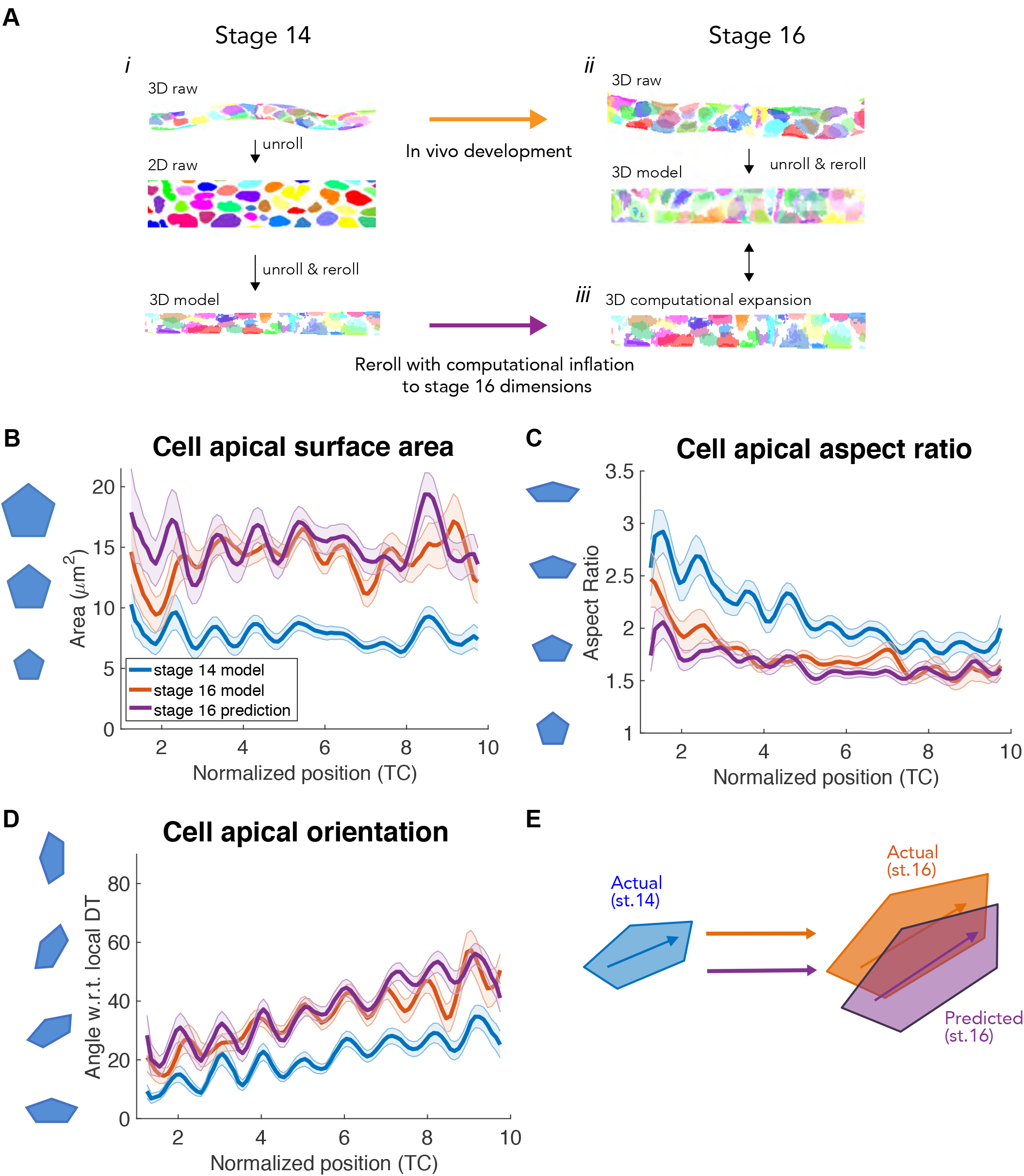
Passive expansion of tracheal cells can account changes in cell apical surface area, aspect ratio and orientation during tube expansion. We used a simple tube expansion paradigm to investigate changes in cell size, shape, and orientation as the trachea grows from stage 14 to stage 16. In outline, we compared data from stage 16 tubes (Aii, orange cell in E) that had undergone in vivo development (orange arrow) to data from stage 14 tubes (Ai, blue cell in E) that had been computational expanded to stage 16 dimensions (Aiii, purple arrow; purple cell in E). **(A) *i.)*** Representative cell apical surfaces from a stage 14 tube (3D raw) and an unrolled tube resulting in a flat 2D image (2D raw). Rerolling results in a 3D model used as a control and for computational inflation (3D model; see text for details). ***ii.)*** Representative cells from a stage 16 tube (3D raw) that had been unrolled and rerolled (3D model). ***iii.)*** Cells on a stage 14 tube that has been computationally expanded to stage 16 (see text for details). **(B-D)** Comparison of cell apical surfaces area, apical aspect ratio and cell orientation between the stage 14 and 16 3D models, and the computational expanded stage 14 3D model. For all parameters, the in vivo stage 16 data (orange lines) show close agreement with the computational expanded stage 14 3D model (purple lines). **(E)** A graphical representation of B-D. During tracheal tube expansion, cells in stage 14 tubes (blue cell) become oriented more orthogonally to the long axis of the tube by stage 16 (orange cell). A similar change in cell angle results from computationally expansion of stage 14 tubes to stage 16 tubes (purple cell), indicating that passive expansion of stage 14 tracheal cells could account for the observed cell geometries at stage 16. Dark lines, mean; shaded path under lines, standard error of the mean.

The cell parameters for stage 16 computationally expanded tubes are shown as purple lines in Figs. 3B-D, with stage 14 and stage 16 models shown as blue and orange lines respectively. As an internal control, we first examined cell apical area (Fig. 3B). Apical area in the computationally expanded tube and actual stage 16 data were comparable, (p = 0.15, paired t-test), with the computational model still showing the strong segmental periodicity observed in the actual data. Interestingly, cell aspect ratio (Fig. 3C) and orientation (Fig. 3D) were also comparable between in vivo stage 16 data and computationally expanded tubes (p = 0.09 and 0.31 respectively, paired t-test), indicating that changes in cell apical shape during tracheal growth are consistent with tracheal cells passively expanding their apical surfaces in response to inflation of the lumen.

### The distribution of dorsal trunk cell connectivity does not change during tube expansion

If tracheal cells simply expanded their apical surface areas in response to lumen inflation, in the simplest case there would be no changes in the cell organization, and the connections between cells would remain static. However, previous work has demonstrated that cell intercalation occurs during tracheal DT growth (Forster and Luschnig, 2012). To test whether QuBiT can find evidence of such intercalations, and to investigate the possible effects of such intercalation, we used QuBiT to measure two quantifiable parameters of cell organization: the number of connections each tracheal cell has with neighboring cells (cell connectivity) and the number of cells per tube cross section.

Cell connectivity can refer to both the number of cells that a given cell shares contacts with, as well as the specific connections a cell has with neighboring cells. Because we analyze data from discrete timepoints rather than time lapse images, we focus on the number of contacts that tracheal cells have, and how that parameter changes along the length of the dorsal trunk and during development. Changes in the distribution in the number of contacts cells have is indicative of migration or other rearrangement events. However, as detailed below, an unchanging connectivity distribution does not demonstrate an absence of cell rearrangement.

To quantify cell connectivity in the trachea, we used QuBiT′s capabilities to unroll the tube to create 2D representations of tracheal cells. We then quantified and visualized cell-cell connectivity using watershed segmentation (Fig. 4A, degrees of connectivity coded by color). Stage 14 and 16 tracheae are predominantly composed of a mix of pentagons and hexagons (Fig. 4B, blue and green bars). There was no obvious clustering of any particular type of polygon at either stage, and the distribution of polygon types is quite uniform along the length of the tube.

**Figure 4.**
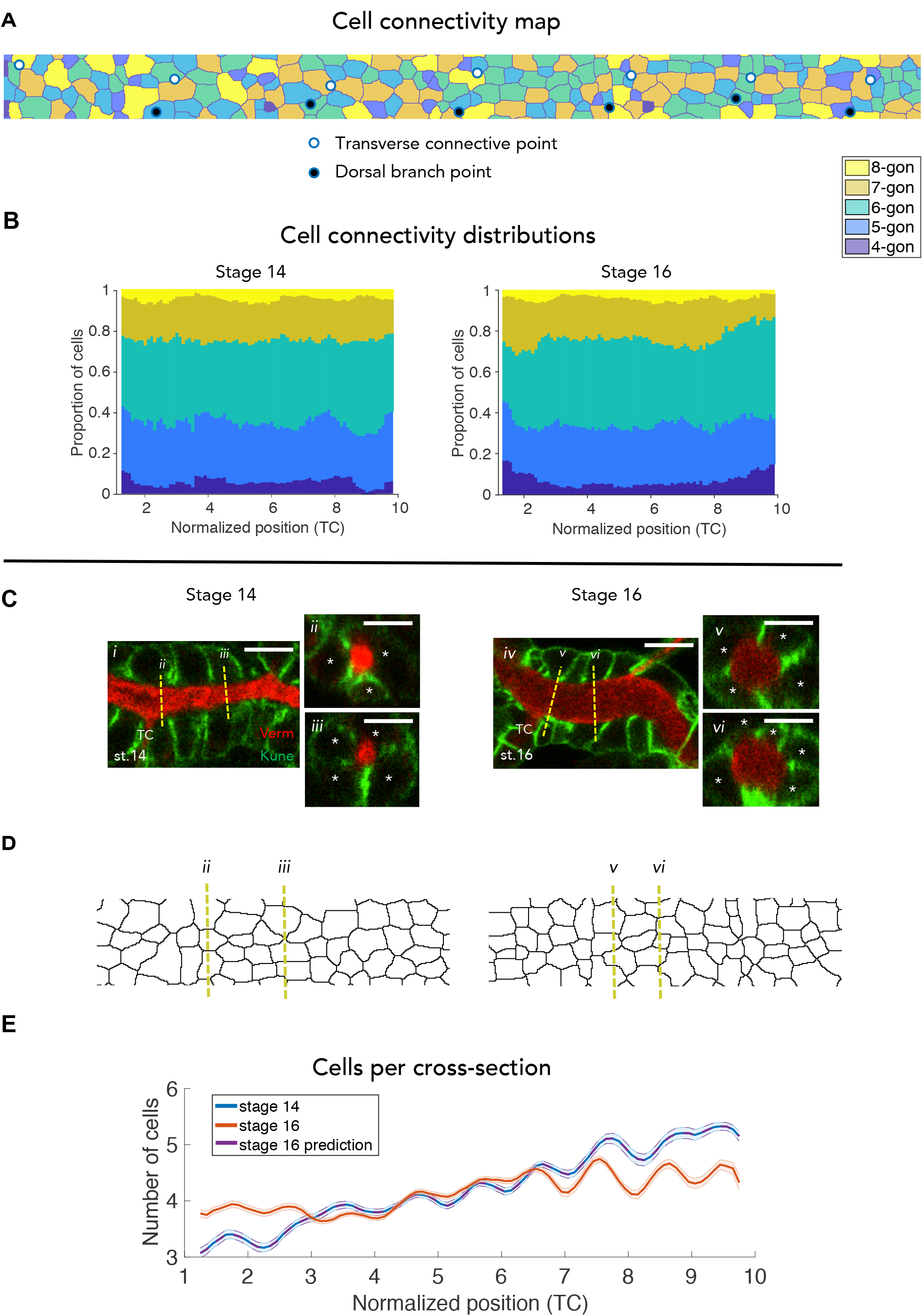
The numbers of cells per cross-section, but not cell connectivity distributions, change during tracheal tube expansion. **(A)** A representative map of tracheal cell connectivity made by unrolling the tube and using watershed segmentation to generate the outlines of cells in 2D. No obvious patterns of connectivity are evident, even at points where the transverse connective and dorsal branches exit the dorsal trunk. X-axis, tube length starting from anterior; y-axis, relative circumferential angular position. Color key, the number of sides they share with neighbors (connectivity). Black and white dots represent DB and TC branch points respectively. **(B)** Bar plots of the distributions of cell connectivity along the DT axis. Cell connectivity distributions do not change from stage 14 to 16. (p = 0.23, χ^2^ test). Color key is same as in A. **(C)** Representative raw images of a single z-slice and their cross-sections (dashed yellow lines) at stages 14 *(i-iii)* and 16 *(iv-vi)* that correspond to the unrolled maps in (D). Scale bar for all images, 5μm. **(D)** 2D unrolled cell maps of the raw images and cross-sections indicated by the dashed yellow lines in (C). **(E)** The number of cells per cross-section changes between stage 14 (blue line) and 16 (orange line) tracheae, indicating cell intercalation events are occurring. The change in the number of cells per crosssection is not predicted by the model of the tracheal cell apical surface passively expanding in response to lumenal inflation (purple line). Interestingly, the number of cells per cross-section becomes more uniform as the tube develops. Also note the presence of a sinusoidal pattern of cells per cross-section superimposed on an A-P gradient, with fewer cells per cross-section near segmental boundaries than in the middle of each tracheal segment.

### The distribution of dorsal trunk cell connectivity is not impacted by side branches

We also took advantage of QuBiT′s ability to accurately map tube branch points to ask whether there were any particular patterns of connectivity associated with the points where the side branches of the DB or TC exit the dorsal trunk (black and white circles in Fig. 4A respectively). No obvious patterns of cell connectivity, or organization of surrounding cell connectivity were observed in 2D maps of tracheal tubes, indicating that neither the complex shapes of the individual cells that contribute to both the dorsal trunk and the side branches nor any forces resulting from tension on the side branches significantly impact the connectivity surrounding dorsal trunk cells.

Importantly, there was no significant change in the distribution of polygons between stages 14 and 16 (Fig. 4B, p = 0.23, χ^2^ test). Thus, it appears that there is little or no change in cell connectivity during tube expansion. This result is consistent with intercalation events occurring infrequently enough that they cause little change to the overall connectivity, but is also consistent with the occurrence of intercalation events conserving their cell connectivity, such as occurs in the T1-T2-T3 transitions and rosette formation in epithelial sheets that drive cell intercalation during convergent extension in zebrafish (Sepich et al., 2000; Tada and Heisenberg, 2012), flies (Blankenship et al., 2006; Irvine and Wieschaus, 1994; Munjal et al., 2015; Pare et al., 2014; Simoes Sde et al., 2014; Zallen and Wieschaus, 2004), and vertebrates (Lienkamp et al., 2012). We further investigated these two possibilities below.

### The number of cells per cross-section changes during tube expansion

To distinguish between a paucity of cell rearrangements versus active changes in cell organization that conserve cell connectivity, we determined the number of cells per cross section of the tube. Infrequent cell intercalations should not significantly change the number of cells per cross section, but in a tube, even intercalation events that conserve cell connectivity would lead to changes in the number of cells per tube cross section if they occurred with significant frequency. We used our 2D diagrams to examine the number of cells in orthogonal cross-sections of the dorsal trunk at stages 14 and 16 (Fig. 4C-D), which yields two notable observations.

First, similar to the patterns observed for cell aspect ratio, orientation and area, a sinusoidal pattern of the number of dorsal trunk cells per cross section also emerges (Fig. 4E). Second, in contrast to the lack of change in connectivity distributions during development, the number of cells per cross section in the dorsal trunk changes from stage 14 to 16, most notably in the anterior and posterior (p < 10^−3^ from TC1 to TC4 and p < 10^−10^ from TC7 to TC10, paired t-test). Critically, since our simple tube expansion model preserves the organization of tracheal cells, the number of cells per cross-section in the computationally expanded stage 16 tube is the same as the original stage 14 tube and is also statistically different from measured stage 16 tubes. Thus, while a simple apical expansion model was sufficient to account for the changes in cell parameters during dorsal trunk expansion, our analytical software indicates cell intercalation events are actually common during tracheal maturation. QuBiT′s ability to predict the existence of cell intercalation events demonstrates that QuBiT can be used to reveal developmental changes in cell organization even in circumstances where live imaging of tube morphogenesis is not feasible.

## Concluding remarks

We created QuBiT to quantify epithelial tube and cell properties in 3D, and to unroll tubes into 2D projections to enable analyses that are either difficult or impossible to do in 3D. The power of this approach is demonstrated by the fact that although the Drosophila tracheal system is one of the most studied epithelial tube systems, QuBiT revealed novel findings. Given these unexpected observations, it is likely that there is much to be found by using QuBiT to analyze tubes in other systems.

## Acknowledgements

We thank Rob Ward for anti-Uif anti-sera, Richard Carthew for helpful discussions, and J. Hornick and the Northwestern University Biological Imaging Facility for imaging support. This work was supported by NIH RO1GM108964 to GJB. M.M. would like to thank the Simons Foundation MMLS program for support.

## Author contributions

R.Y., M.M. and G.J.B conceived and designed the study, R.Y., E.L., M.M. and G.J.B developed the method. R.Y., M.M. and G.J.B analyzed the data and wrote the paper. All of the authors approved the manuscript.

---

**Figure S1.**
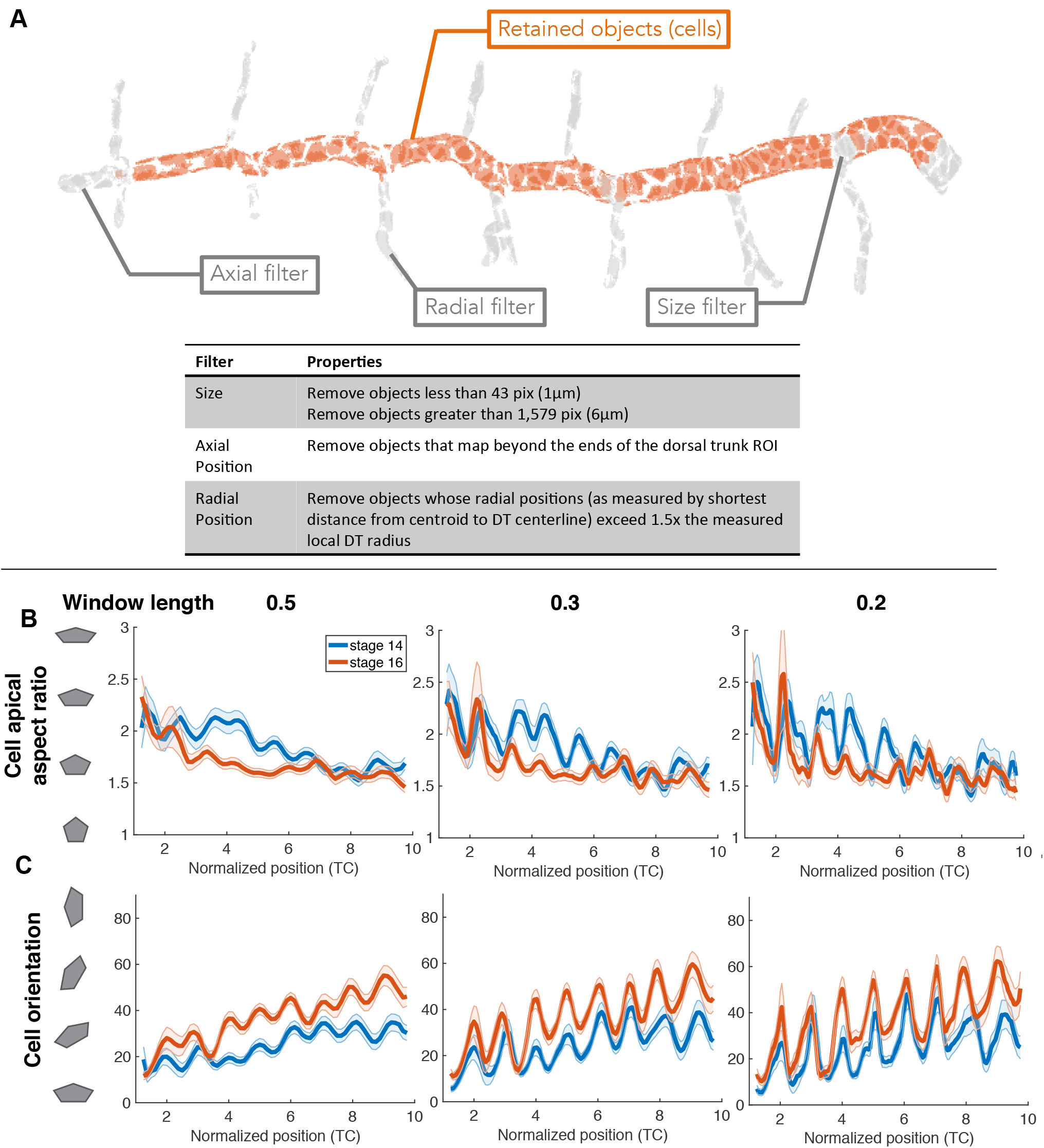
Filtering and assessment of effect of window size on segmental patterns. **(A)** Visual representation of filtering to identify unique cells. We applied three different filters to remove incorrectly segmented objects (grey cells). The axial filter removed cells at the ends of the tube that were out of range of our dorsal trunk region of interest. The radial filter removed cells that exceeded 1.5× the measured local radius, which corresponded to approximately 1.5μm (anterior) to 2.5μm (posterior) in stage 14 tubes and 3μm to 5μm in stage 16 tubes. The size filter removed objects that were improbably small or large (<1 μm and >6 μm respectively), based on empirical observations of conjoined cells due to poor segmentation and objects which had sizes beyond 3 standard deviations from the mean. The remaining objects are considered cells and are labeled in orange. **(B-C)** Tracheal cell apical aspect ratio and orientation show segmental organization as well as A-P gradients. To demonstrate that the periodicity in aspect ratio and orientation does not result from an artifact of the size of the window used for averaging along the A-P tube axis, we varied window size from 50% to 20% of the length of a segment. The periodicity and A-P gradients are evident in all cases.

**Figure S2.**
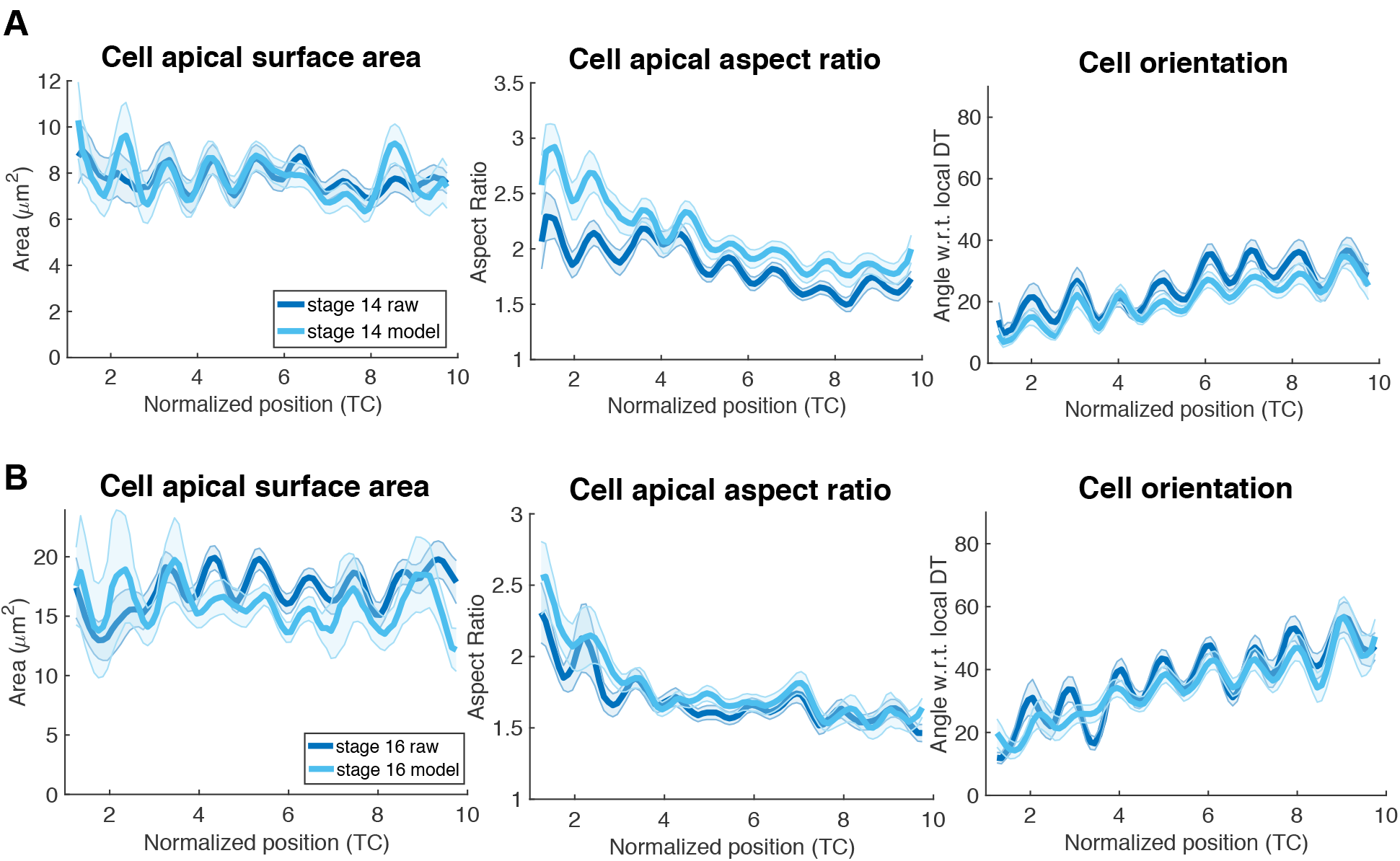
Assessment of the effects of tube straightening. **(A-B)** Controls for computational expansion of tracheal tubes. The workflow for computationally expanding tracheal tube is shown in Fig. 3A. As detailed in the text, during the expansion, the tube is unrolled and then rerolled and expanded onto straight long axis, which causes some deformations (note alternative approaches also cause deformation). To verify that straightening (modeling) the tube maintains cell shapes similar to their original shapes, we plotted the straightened cell properties (surface area, aspect ratio, and orientation) against the data from native tubes at (A) stage 14 and (B) stage 16. Most parameters are in excellent agreement. However, cell apical aspect ratios at stage 14 differ between our raw and model data. This is most likely due to the narrow tube diameter relative to cell size, which magnifies small deviations in cell shape (note that the inconsistency is more apparent at the anterior, where tubes are thinnest).

